## Supplemental Tables, FIgures, Methods for "Previously uncharacterized rectangular bacterial structures in the dolphin mouth"

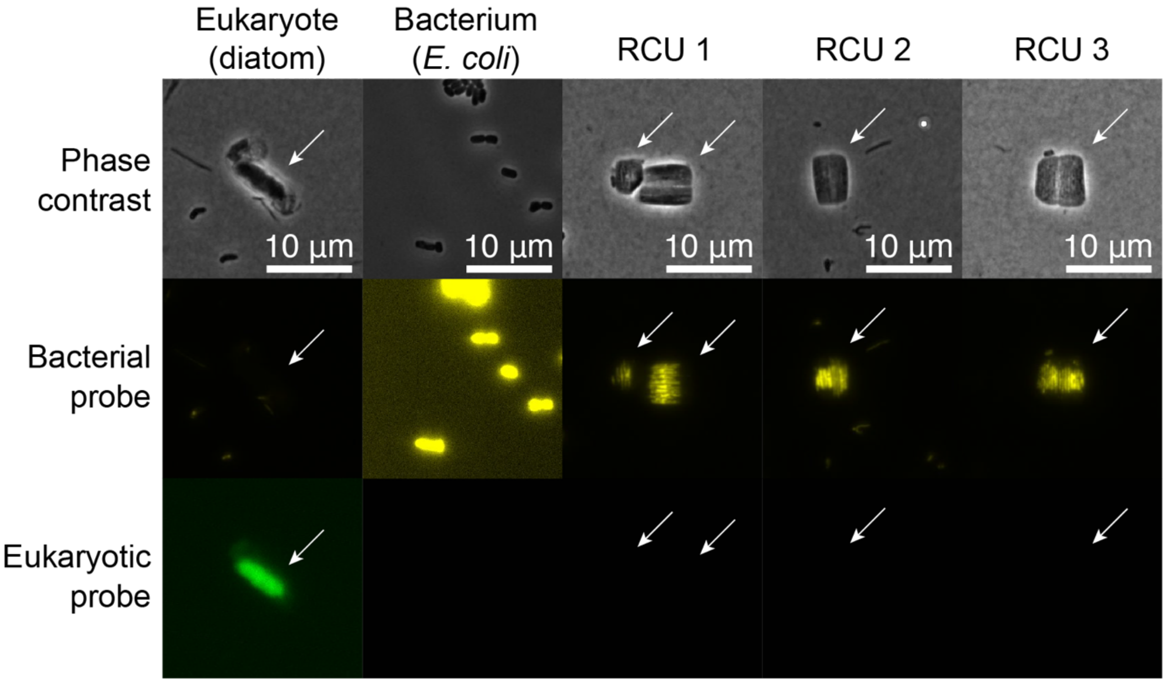


**Supplementary Figure 1. Fluorescence *in situ* hybridization indicates that RBSs are bacterial rather than eukaryotic.** Bacterial probe Eub-338 was labeled with AlexaFluor-488 and eukaryotic probe Euk-1209 was labeled with AlexaFluor-660. Top: phase-contrast images; middle, bottom: fluorescence images from bacterial and eukaryotic probes, respectively. Arrows indicate the relevant cells in non-axenic samples. The first column is a cell of the marine diatom *Skeletonema costatum* grown in non-axenic culture. The second column is axenic *Escherichia coli* cells. The last three columns are RBSs obtained directly from dolphin oral swabs. The bacterial probe labeled all RBSs, while the eukaryotic probe only labeled the marine diatom.

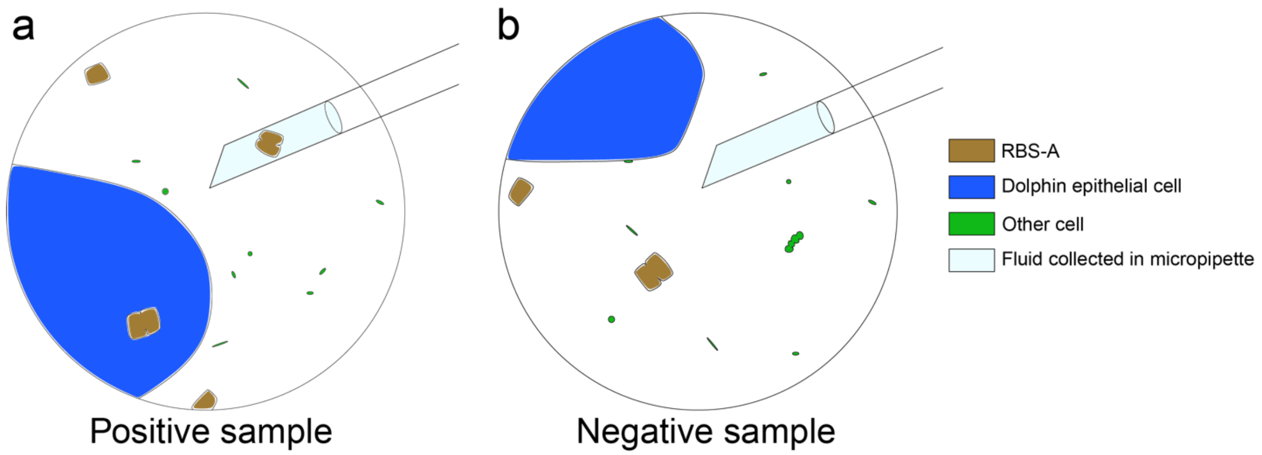


**Supplementary Figure 2. Collection strategy for single-cell genomics.** Four samples of RBS-As were collected, each with ~1-3 RBS-As per tube. Four negative control samples with fluid from the dolphin oral sample but not RBS-As were also collected. While care was taken to avoid collecting non-RBS-A cells, it is likely that small, non-visualized cells and/or cell-free DNA were captured along with RBS-As.

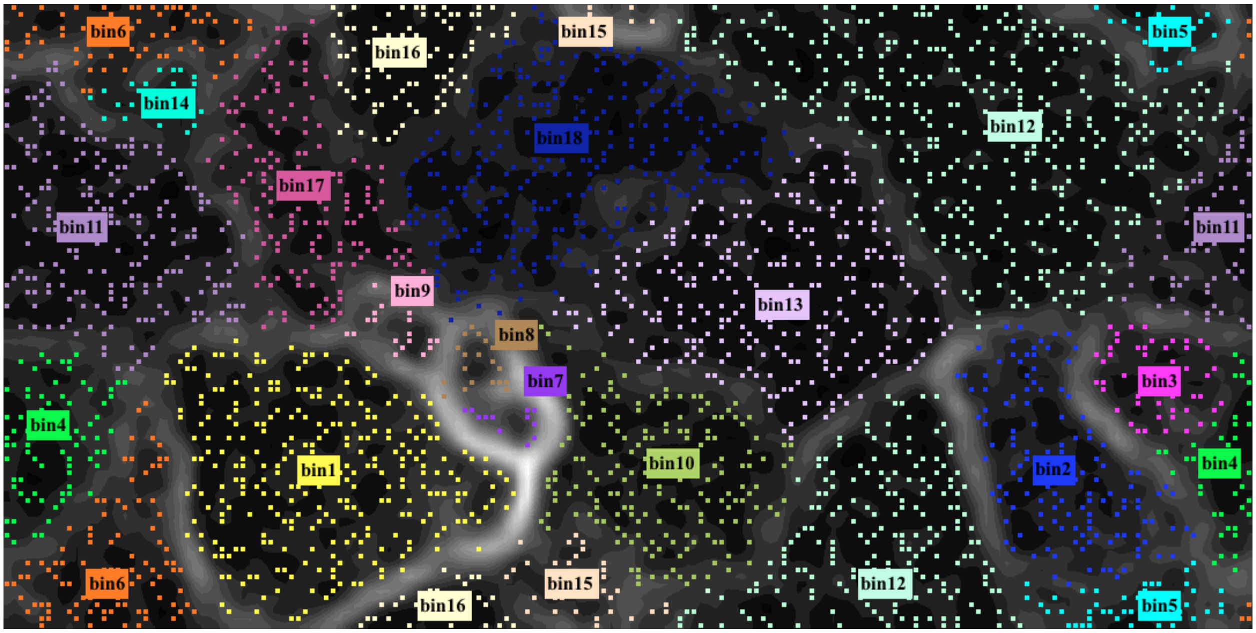


**Supplementary Figure 3. Tetranucleotide Emergent Self-Organizing Map (ESOM) used to bin genomes.** Scaffolds were split into keys of size 5000. Keys (dots on the ESOM), each of which represents a 5-kb section of scaffold, are color-coded based on the bin to which they were assigned.

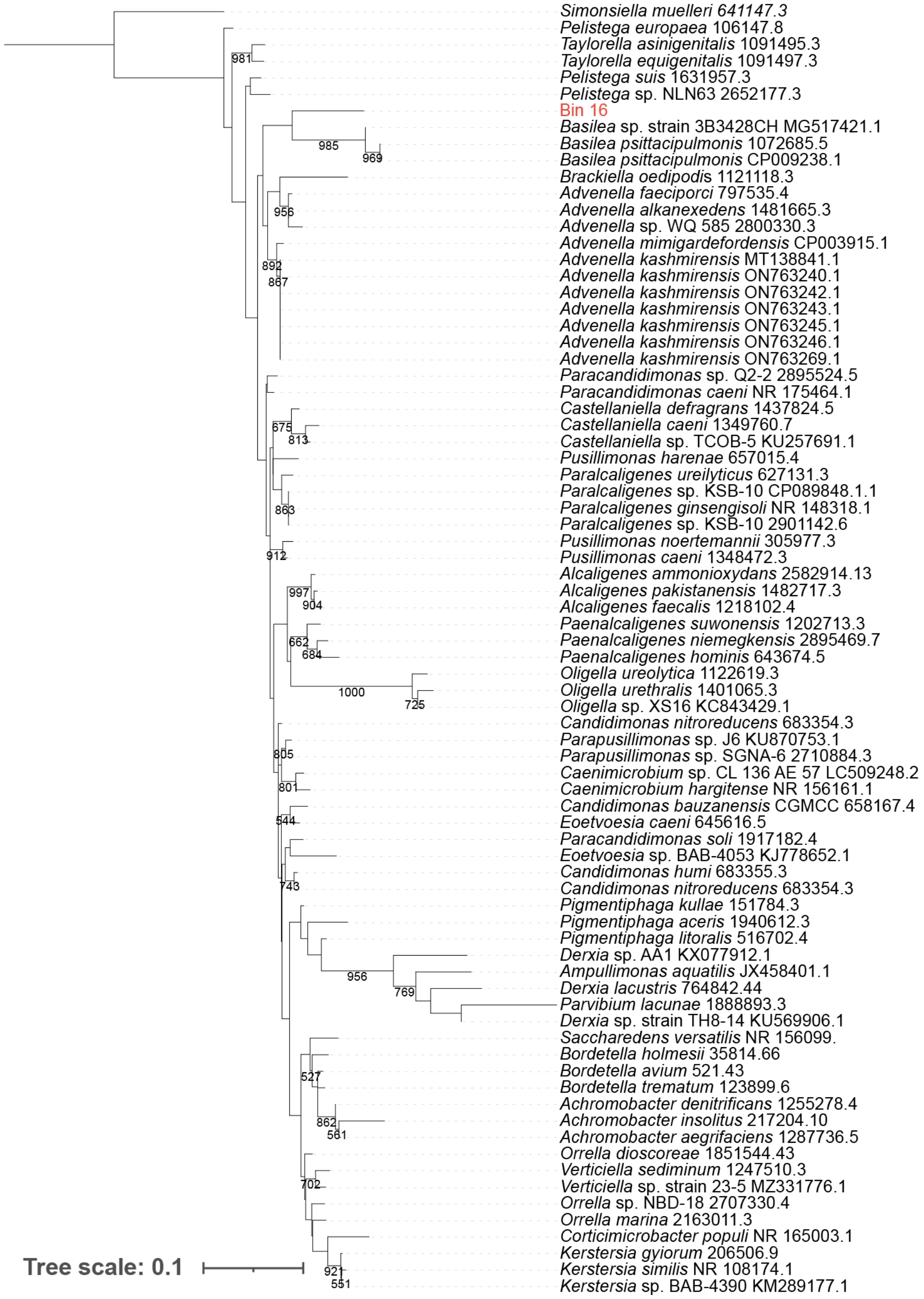


**Supplementary Figure 4. Phylogenetic analysis of the 16S rRNA gene confirms that bin 16 is affiliated with the Alcaligenaceae.** Maximum likelihood 16S rRNA gene phylogeny of the Alcaligenaceae with 1000 bootstraps. Up to three representative sequences from each Alcaligenaceae genus, as listed on the NCBI Taxonomy Browser, were included, as were the top 10 gene sequences most similar to the bin 16 16S rRNA gene as identified through a BLAST[1] query against the NCBI nr/nt database. Bin 16 is highlighted in red; other leaves are represented by species name and NCBI identifier.

Bootstrap support values ≥50% (≥500) are shown. The 16S rRNA gene of *S. muelleri* was used as an outgroup.

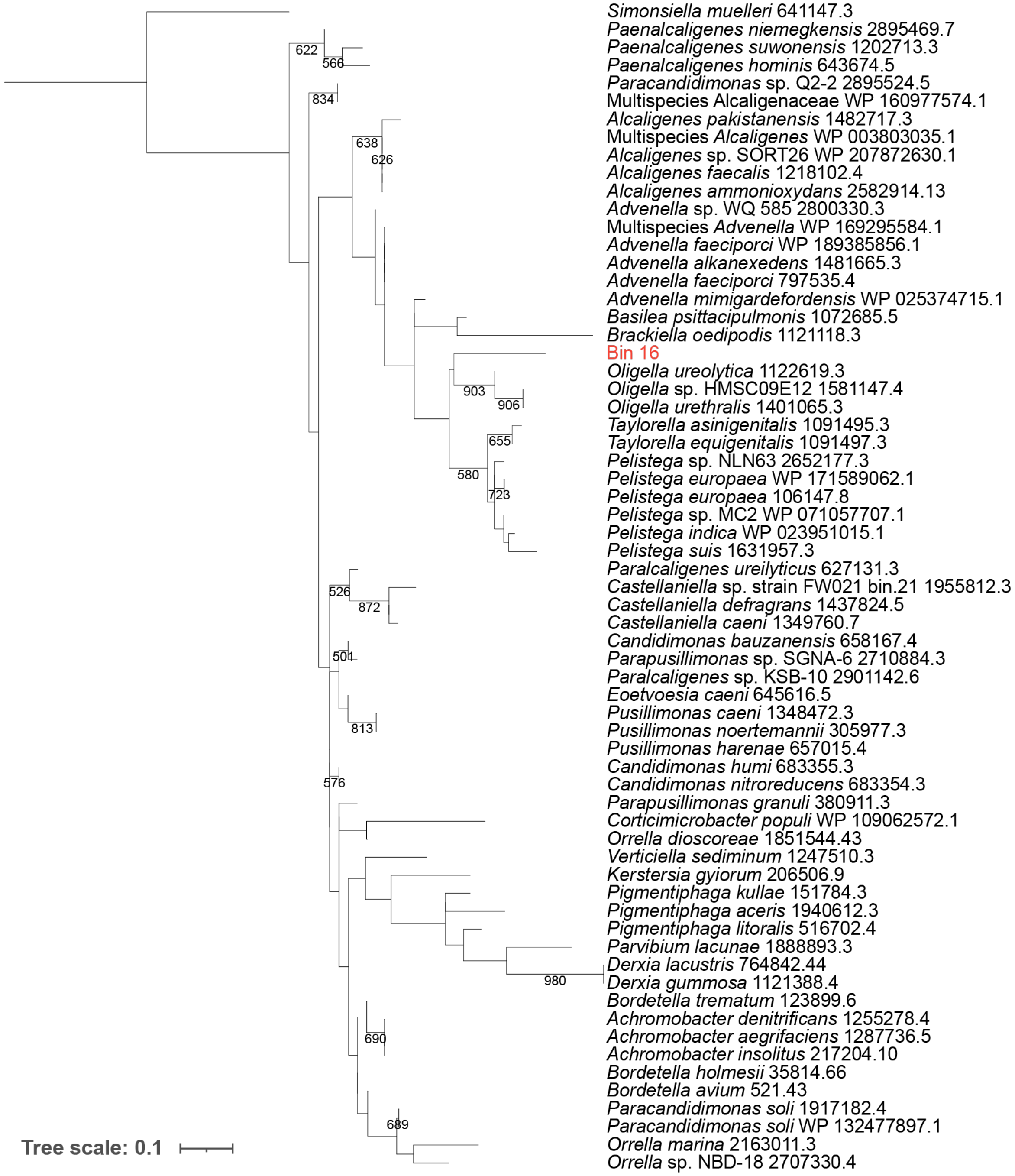


**Supplementary Figure 5. Phylogenetic analysis of ribosomal protein S3 (rpS3) confirms bin 16 is affiliated with the Alcaligenaceae.** Maximum likelihood rpS3 phylogeny of the Alcaligenaceae with 1000 bootstraps. Up to three representative sequences from each Alcaligenaceae genus, as listed on the NCBI Taxonomy Browser, were included, as were the top 10 protein sequences most similar to the bin 16 rpS3 as identified through a BLAST[1] query against the NCBI nr database. Bin 16 is highlighted in red; other leaves are represented by species name and NCBI identifier. Bootstrap support values ≥50% (≥500) are shown. The rpS3 protein sequence of *S. muelleri* was used as an outgroup.

| Sample location | FOVs | Dolphin | Collection Date | Morphotype A | Morphotype B | Amplicon sequencing |
|---|---|---|---|---|---|---|
| Palate, tongue, gingivus | 226 | 1 | Jan-15-2018 | *** | 0 | Y |
| Palate, tongue, gingivus | 226 | 1 | Jan-15-2018 | ** | * | Y |
| Palate, tongue, gingivus | 226 | 1 | Jan-18-2018 | 0 | 0 | Y |
| Palate, tongue, gingivus | 226 | 1 | Jan-18-2018 | 0 | 0 | Y |
| Palate, tongue, gingivus | 226 | 1 | Jan-22-2018 | 0 | 0 | Y |
| Palate, tongue, gingivus | 226 | 1 | Jan-22-2018 | 0 | 0 | Y |
| Palate, tongue, gingivus | 226 | 1 | Jan-22-2018 | 0 | 0 | Y |
| Palate, tongue, gingivus | 226 | 1 | Jan-25-2018 | 0 | 0 | Y |
| Palate, tongue, gingivus | 226 | 1 | Jan-25-2018 | 0 | 0 | Y |

| | | | | | | |
|---|---|---|---|---|---|---|
| Palate, tongue, gingivus | 226 | 2 | Jan-15-2018 | 0 | ** | Y |
| Palate, tongue, gingivus | 226 | 2 | Jan-18-2018 | ** | * | Y |
| Palate, tongue, gingivus | 226 | 2 | Jan-18-2018 | ** | ** | Y |
| Palate, tongue, gingivus | 226 | 2 | Jan-22-2018 | * | ** | Y |
| Palate, tongue, gingivus | 226 | 2 | Jan-22-2018 | 0 | ** | Y |
| Palate, tongue, gingivus | 226 | 2 | Jan-25-2018 | 0 | ** | Y |
| Palate, tongue, gingivus | 226 | 2 | Jan-25-2018 | 0 | ** | Y |
| Palate, tongue, gingivus | 226 | 3 | Jan-29-2018 | ** | * | Y |
| Palate, tongue, gingivus | 226 | 3 | Jan-29-2018 | ** | * | Y |
| Palate, tongue, gingivus | 226 | 3 | Feb-1-2018 | ** | 0 | Y |

| | | | | | | |
|---|---|---|---|---|---|---|
| Palate, tongue, gingivus | 226 | 3 | Feb-1-2018 | * | 0 | Y |
| Palate, tongue, gingivus | 226 | 3 | Feb-8-2018 | * | * | Y |
| Palate, tongue, gingivus | 226 | 3 | Feb-8-2018 | * | 0 | Y |
| Palate, tongue, gingivus | 226 | 4 | Jan-29-2018 | ** | 0 | Y |
| Palate, tongue, gingivus | 226 | 4 | Feb-1-2018 | 0 | 0 | Y |
| Palate, tongue, gingivus | 226 | 4 | Feb-8-2018 | * | 0 | Y |
| Palate, tongue, gingivus | 226 | 4 | Feb-8-2018 | 0 | 0 | Y |
| Palate, tongue, gingivus | 226 | 4 | Feb-8-2018 | 0 | 0 | Y |
| Palate, tongue, gingivus | 226 | 5 | Feb-12-2018 | *** | ** | Y |
| Palate, tongue, gingivus | 226 | 5 | Feb-15-2018 | ** | ** | Y |

| | | | | | | |
|---|---|---|---|---|---|---|
| Palate, tongue, gingivus | 226 | 5 | Feb-15-2018 | 0 | * | Y |
| Palate, tongue, gingivus | 226 | 5 | Feb-19-2018 | 0 | 0 | Y |
| Palate, tongue, gingivus | 226 | 5 | Feb-19-2018 | 0 | 0 | Y |
| Palate, tongue, gingivus | 226 | 5 | Feb-22-2018 | 0 | 0 | Y |
| Palate, tongue, gingivus | 226 | 5 | Feb-22-2018 | 0 | 0 | Y |
| Palate, tongue, gingivus | 226 | 5 | Feb-22-2018 | 0 | 0 | Y |
| Palate, tongue, gingivus | 226 | 6 | Feb-12-2018 | ** | * | Y |
| Palate, tongue, gingivus | 226 | 6 | Feb-12-2018 | * | * | Y |
| Palate, tongue, gingivus | 226 | 6 | Feb-15-2018 | ** | 0 | Y |
| Palate, tongue, gingivus | 226 | 6 | Feb-15-2018 | ** | * | Y |

| | | | | | | |
|---|---|---|---|---|---|---|
| Palate, tongue, gingivus | 226 | 6 | Feb-15-2018 | * | 0 | Y |
| Palate, tongue, gingivus | 226 | 6 | Feb-19-2018 | * | ** | Y |
| Palate, tongue, gingivus | 226 | 6 | Feb-19-2018 | 0 | ** | Y |
| Palate, tongue, gingivus | 226 | 6 | Feb-22-2018 | 0 | ** | Y |
| Palate, tongue, gingivus | 226 | 6 | Feb-22-2018 | 0 | * | Y |
| Gingivus | 226 | 7 | Apr-1-2012 | * | * | Y |
| Palate, tongue, gingivus | 226 | 8 | Jan-25-2018 | 0 | 0 | Y |
| Palate, tongue, gingivus | 226 | 8 | Jan-25-2018 | 0 | 0 | Y |
| Palate, tongue, gingivus | 226 | 8 | Jan-25-2018 | 0 | 0 | Y |
| Buccal surface | 100 | 3 | Mar-24-2022 | 0 | 0 | N |
| Buccal surface | 100 | 4 | Mar-24-2022 | 0 | 0 | N |
| Buccal surface | 100 | 5 | Mar-24-2022 | 0 | 0 | N |

| | | | | | | |
|---|---|---|---|---|---|---|
| Buccal surface | 100 | 6 | Mar-24-2022 | 0 | * | N |
| Buccal surface | 100 | 8 | Mar-24-2022 | 0 | * | N |
| Gingivus | 100 | 5 | Mar-24-2022 | *** | ** | N |
| Gingivus | 100 | 7 | Mar-24-2022 | *** | ** | N |
| Gingivus | 100 | 5 | Mar-24-2022 | ** | * | N |
| Gingivus | 100 | 6 | Mar-24-2022 | ** | ** | N |
| Gingivus | 100 | 7 | Mar-24-2022 | ** | ** | N |
| Gingivus | 100 | 3 | Mar-24-2022 | * | 0 | N |
| Gingivus | 100 | 3 | Mar-24-2022 | * | * | N |
| Gingivus | 100 | 3 | Mar-24-2022 | * | * | N |
| Gingivus | 100 | 5 | Mar-24-2022 | * | * | N |
| Gingivus | 100 | 6 | Mar-24-2022 | * | * | N |
| Gingivus | 100 | 4 | Mar-24-2022 | * | ** | N |
| Gingivus | 100 | 6 | Mar-24-2022 | * | ** | N |
| Gingivus | 100 | 4 | Mar-24-2022 | 0 | * | N |

| | | | | | | |
|---|---|---|---|---|---|---|
| Gingivus | 100 | 4 | Mar-24-2022 | 0 | * | N |
| Gingivus | 100 | 7 | Mar-24-2022 | 0 | *** | N |
| Palate | 100 | 6 | Mar-24-2022 | *** | 0 | N |
| Palate | 100 | 3 | Mar-24-2022 | *** | * | N |
| Palate | 100 | 5 | Mar-24-2022 | ** | * | N |
| Palate | 100 | 4 | Mar-24-2022 | ** | ** | N |
| Palate | 100 | 7 | Mar-24-2022 | * | ** | N |

**Supplementary Table 1.** Prevalence of RBSs in dolphin oral samples. 73 samples from eight dolphins were evaluated for the presence of RBSs using SLIP[2] via phase-contrast microscopy. The order of magnitude of RBS counts for each morphology per sample is shown here; * denotes 1-9 RBSs of a given morphotype detected, ** denotes 10-99, and *** denotes ≥100 (maximum identified in a single sample was 366). Samples collected prior to 2022 were surveyed using 226 fields of view (FOVs) and underwent amplicon sequencing, while those collected in 2022 were surveyed using 100 FOVs and did not undergo sequencing. RBS-As and RBS-Bs were each detected in 39/73 samples (53%) and 42/73 samples (58%), respectively. RBS-As were detected in samples from 7/8 dolphins, while RBS-Bs were detected in samples from 6/8 dolphins.

| ASV | # samples | Phylum | Class | Order | Family | Genus | Species |
|---|---|---|---|---|---|---|---|
| 1 | 13 | Proteobacteria | Gammaproteobacteria | Pasteurellales | Pasteurellaceae | *Pasteurella* | *skyensis* |
| 2 | 13 | Proteobacteria | Gammaproteobacteria | Pseudomonadales | Moraxellaceae | - | - |
| 3 | 13 | Proteobacteria | Gammaproteobacteria | Cardiobacteriales | - | - | - |
| 4 | 13 | Proteobacteria | Gammaproteobacteria | Pseudomonadales | Moraxellaceae | - | - |
| 5 | 13 | Bacteroidetes | Flavobacteriia | Flavobacteriales | Flavobacteriaceae | - | - |
| 6 | 13 | Fusobacteria | Fusobacteriia | Fusobacteriales | Fusobacteriaceae | *Fusobacterium* | - |
| 7 | 13 | Proteobacteria | Gammaproteobacteria | Pasteurellales | Pasteurellaceae | *Pasteurella* | *skyensis* |
| 8 | 13 | Proteobacteria | Epsilonproteobacteria | Campylobacterales | Campylobacteraceae | *Arcobacter* | - |
| 9 | 13 | Proteobacteria | Epsilonproteobacteria | Campylobacterales | Campylobacteraceae | - | - |
| 10 | 13 | Firmicutes | Bacilli | Lactobacillales | Enterococcaceae | *Enterococcus* | - |
| 11 | 12 | Proteobacteria | Gammaproteobacteria | Cardiobacteriales | - | - | - |
| 12 | 12 | Bacteroidetes | Flavobacteriia | Flavobacteriales | Flavobacteriaceae | *Tenacibaculum* | - |
| 13 | 12 | Bacteroidetes | - | - | - | - | - |
| 14 | 12 | Bacteroidetes | - | - | - | - | - |
| 15 | 12 | Bacteroidetes | Flavobacteriia | Flavobacteriales | Flavobacteriaceae | *Tenacibaculum* | - |
| 16 | 11 | Bacteroidetes | Bacteroidia | Bacteroidales | Porphyromonadaceae | *Porphyromonas* | - |
| 17 | 11 | Proteobacteria | Gammaproteobacteria | Cardiobacteriales | - | - | - |
| 18 | 11 | Bacteroidetes | Bacteroidia | Bacteroidales | Porphyromonadaceae | *Paludibacter* | - |
| 19 | 11 | Proteobacteria | Betaproteobacteria | Burkholderiales | Alcaligenaceae | - | - |
| 20 | 10 | Proteobacteria | Gammaproteobacteria | Pseudomonadales | Moraxellaceae | - | - |

| | | | | | | | |
|---|---|---|---|---|---|---|---|
| 21 | 10 | Proteobacteria | Gammaproteobacteria | Cardiobacteriales | - | - | - |
| 22 | 10 | Bacteroidetes | Flavobacteriia | Flavobacteriales | Flavobacteriaceae | *Tenacibaculum* | - |
| 23 | 10 | GN02 | BD1-5 | - | - | - | - |
| 24 | 10 | Firmicutes | Clostridia | Clostridiales | Lachnospiraceae | - | - |

**Supplemental Table 2. Taxonomic ID of ASVs present in ≥75% of samples with RBS-As detected.** Only samples with ≥10 RBS-As visually confirmed were considered to bolster confidence in the assessment (e.g., that an RBS-B was not accidentally considered as an RBS-A). Out of 13 samples meeting this criterion, the number of samples in which any given ASV appears is shown, along with assigned taxonomic ID at the lowest possible taxonomic level. Dashes ("-") indicate that an ASV was not assigned at a given taxonomic level. Note that the Alcaligenaceae ASV has 100% sequence identity over 100% length to the 16S rRNA gene associated with the Alcaligenaceae bin recovered from the mini-metagenomics experiment.

| ASV | # samples | Phylum | Class | Order | Family | Genus | Species |
|---|---|---|---|---|---|---|---|
| 1 | 11 | Proteobacteria | Gammaproteobacteria | Pasteurellales | Pasteurellaceae | *Pasteurella* | *skyensis* |
| 2 | 11 | Proteobacteria | Gammaproteobacteria | Pseudomonadales | Moraxellaceae | - | - |
| 3 | 11 | Proteobacteria | Gammaproteobacteria | Cardiobacteriales | - | - | - |
| 4 | 11 | Bacteroidetes | Flavobacteriia | Flavobacteriales | Flavobacteriaceae | - | - |
| 5 | 11 | Fusobacteria | Fusobacteriia | Fusobacteriales | Fusobacteriaceae | *Fusobacterium* | - |
| 6 | 11 | Proteobacteria | Gammaproteobacteria | Pasteurellales | Pasteurellaceae | *Pasteurella* | *skyensis* |
| 7 | 11 | Bacteroidetes | Flavobacteriia | Flavobacteriales | Flavobacteriaceae | *Tenacibaculum* | - |
| 8 | 11 | Firmicutes | Bacilli | Lactobacillales | Enterococcaceae | *Enterococcus* | - |
| 9 | 10 | Proteobacteria | Gammaproteobacteria | Pseudomonadales | Moraxellaceae | - | - |
| 10 | 10 | Proteobacteria | Gammaproteobacteria | Cardiobacteriales | - | - | - |
| 11 | 10 | Bacteroidetes | Bacteroidia | Bacteroidales | - | - | - |
| 12 | 10 | Bacteroidetes | Flavobacteriia | Flavobacteriales | Flavobacteriaceae | - | - |
| 13 | 10 | Bacteroidetes | Bacteroidia | Bacteroidales | Porphyromonadaceae | *Paludibacter* | - |
| 14 | 10 | Bacteroidetes | Flavobacteriia | Flavobacteriales | [Weeksellaceae] | - | - |

| | | | | | | | |
|---|---|---|---|---|---|---|---|
| 15 | 9 | Proteobacteria | Gammaproteobacteria | Pseudomonadales | Moraxellaceae | - | - |
| 16 | 9 | Proteobacteria | Epsilonproteobacteria | Campylobacterales | Campylobacteraceae | *Arcobacter* | - |
| 17 | 9 | Proteobacteria | Gammaproteobacteria | Cardiobacteriales | - | - | - |
| 18 | 9 | Bacteroidetes | Flavobacteriia | Flavobacteriales | Flavobacteriaceae | *Tenacibaculum* | - |
| 19 | 9 | Proteobacteria | Epsilonproteobacteria | Campylobacterales | Campylobacteraceae | - | - |
| 20 | 9 | Bacteroidetes | Flavobacteriia | Flavobacteriales | Flavobacteriaceae | - | - |
| 21 | 9 | Proteobacteria | Betaproteobacteria | Burkholderiales | Alcaligenaceae | - | - |
| 22 | 9 | Proteobacteria | Gammaproteobacteria | Cardiobacteriales | - | - | - |
| 23 | 9 | Proteobacteria | Epsilonproteobacteria | Campylobacterales | Campylobacteraceae | - | - |
| 24 | 9 | Bacteroidetes | Flavobacteriia | Flavobacteriales | Flavobacteriaceae | *Tenacibaculum* | - |
| 25 | 9 | Bacteroidetes | Bacteroidia | Bacteroidales | Bacteroidaceae | *Bacteroides* | *ovatus* |
| 26 | 9 | Firmicutes | Clostridia | Clostridiales | Lachnospiraceae | - | - |

**Supplemental Table 3. Taxonomic ID of ASVs present in ≥75% of samples with RBS-Bs detected.** Only samples with ≥10 RBS-As visually confirmed were considered to bolster confidence in the assessment (e.g., that an RBS-A was not accidentally considered as an RBS-B). Out of 13 samples meeting this criterion, the number of samples in which any given ASV appears is shown, along with the assigned taxonomic ID at the lowest possible taxonomic level. Dashes (“-”) indicate that an ASV was not assigned at a given taxonomic level. Note that the Alcaligenaceae ASV has 100% sequence identity over 100% length to the 16S rRNA gene associated with the Alcaligenaceae bin recovered from the mini-metagenomics experiment.

| Bin ID | # scaffolds | Bin length | Longest scaffold | N50 | Completeness (%) | Contamination (%) |
|---|---|---|---|---|---|---|
| 1 | 272 | 1,941,293 | 26,105 | 6,813 | 1 | 0 |
| 2 | 56 | 846,971 | 68,899 | 18,366 | 49 | 6 |
| 3 | 25 | 288,673 | 27,320 | 10,819 | 14 | 0 |
| 4 | 45 | 732,195 | 52,808 | 24,418 | 34 | 0 |
| 5 | 36 | 434,317 | 31,157 | 15,055 | 18 | 0 |
| 6 | 93 | 1,026,923 | 32,958 | 12,480 | 59 | 0 |
| 7 | 16 | 116,224 | 14,175 | 7,023 | 2 | 0 |
| 8 | 15 | 127,029 | 15,569 | 8,224 | 0 | 0 |
| 9 | 12 | 92,216 | 12,253 | 7,606 | 0 | 0 |
| 10 | 121 | 1,176,422 | 35,727 | 9,462 | 31 | 2 |
| 11 | 56 | 1,591,011 | 224,748 | 40,556 | 76 | 1 |
| 12 | 204 | 3,272,710 | 104,534 | 23,600 | 90 | 40 |
| 13 | 187 | 1,783,018 | 24,881 | 10,073 | 58 | 10 |
| 14 | 12 | 137,833 | 25,347 | 14,892 | 7 | 0 |
| 15 | 21 | 171,018 | 13,722 | 99 | 0 | 0 |
| 16 | 43 | 712,931 | 90,478 | 16,606 | 42 | 1 |
| 17 | 89 | 801,486 | 40,325 | 9,084 | 12 | 0 |
| 18 | 68 | 1,679,601 | 97,578 | 33,569 | 83 | 15 |

**Supplementary Table 4. Assembly statistics for bins recovered from the mini-metagenomics experiment.** For each bin, the number of scaffolds, total length of all scaffolds, longest scaffold, N50, completeness (%), and contamination (%) are reported. Completeness and contamination were estimated based on the presence/absence of a broad set of marker genes using CheckM1.

| | RBS-A samples | | | | Negative controls | | | |
|---|---|---|---|---|---|---|---|---|
| **Bin ID** | **1** | **2** | **3** | **4** | **1** | **2** | **3** | **4** |
| 1 | 0.00 | 0.00 | 0.05 | 0.00 | 0.00 | 0.02 | 0.00 | 87.74 |
| 2 | 8.44 | 1.29 | 0.05 | 16.58 | 0.00 | 61.22 | 0.00 | 0.00 |
| 3 | 0.00 | 0.00 | 0.00 | 4.17 | 0.00 | 0.00 | 0.00 | 0.66 |
| 4 | 0.00 | 1.23 | 0.00 | 4.35 | 0.00 | 0.00 | 0.00 | 0.00 |
| 5 | 10.94 | 29.80 | 0.00 | 0.24 | 0.00 | 2.64 | 0.00 | 0.26 |
| 6 | 2.84 | 0.06 | 0.00 | 3.04 | 0.00 | 0.00 | 0.00 | 1.91 |
| 7 | 0.00 | 0.00 | 0.00 | 0.00 | 0.00 | 6.32 | 0.00 | 6.74 |
| 8 | 0.00 | 0.00 | 0.00 | 0.00 | 0.00 | 8.84 | 0.00 | 0.00 |
| 9 | 1.49 | 0.44 | 0.00 | 2.31 | 92.13 | 13.85 | 0.00 | 0.00 |
| 10 | 15.22 | 9.90 | 0.02 | 15.61 | 0.00 | 1.11 | 0.00 | 1.50 |
| 11 | 1.11 | 0.00 | 0.00 | 11.52 | 0.00 | 0.59 | 0.00 | 0.00 |
| 12 | 14.53 | 12.32 | 3.98 | 3.67 | 3.83 | 2.57 | 0.00 | 0.60 |
| 13 | 16.49 | 3.73 | 47.83 | 1.77 | 4.04 | 0.58 | 0.00 | 0.06 |
| 14 | 0.01 | 0.07 | 0.00 | 2.35 | 0.00 | 0.08 | 0.00 | 0.00 |
| 15 | 9.31 | 37.48 | 1.39 | 5.07 | 0.00 | 0.38 | 0.00 | 0.00 |
| 16 | 13.70 | 0.00 | 0.00 | 0.21 | 0.00 | 0.00 | 0.00 | 0.00 |
| 17 | 3.37 | 3.52 | 45.62 | 6.85 | 0.00 | 1.80 | 0.00 | 0.52 |
| 18 | 2.57 | 0.15 | 1.05 | 22.27 | 0.00 | 0.00 | 0.00 | 0.00 |
| **# read pairs** | 2,751,812 | 2,896,207 | 6,306,907 | 5,051,639 | 2,201,169 | 3,522,055 | 3,197,485 | 5,059,659 |
| **# proper read pair alignments** | 487,042 | 119,453 | 7,013 | 511,482 | 769 | 241,222 | 0 | 185,809 |

**Supplementary Table 5. Relative abundance (%) of bins recovered in the mini-metagenomics experiment.** We applied the following thresholds to aid in the evaluation of whether a given bin was likely to have been present in a sample, color-coded by relative abundance as follows: green, ≥ 5%; yellow, ≥ 1% and < 5%; orange, ≥ 0% and < 1%, red: 0%. The last two rows of the table show the total number of read pairs per sample and the number of primary alignments with proper read pairs mapping to scaffolds ≥ 5 kb long per sample (only scaffolds ≥ 5 kb were binned; see **Methods**).

| TEM instrument | Data collection type | Pixel size (Å/pixel) | Corresponding figure(s) | Applied defocus (μm) | Cumulative dose ($e^{-}$/Å$^{2}$) |
|---|---|---|---|---|---|
| Titan Krios G3, 300 keV, energy filter at 20-eV slit width, K2 detector | Tilt series | 3.48 | Figure 7C | -6 | 120 |
| | 2D montage maps and/or images | 1.06 | Figure 7A<br>Figure S7D | -5 | 80 |
| | | 3.48 | Figure 7B<br>Figure 5D | -5 | |
| Titan Krios G2, 300 keV, no energy filter, K2 detector | Tilt series | 3.75 | Figure 6B | -6 | 120 |
| | | 7.5 | Figure 6A | -6 | |
| | 2D montage maps and/or images | 1.43 | Figure 5A<br>Figure 5E | -5 | 80 |
| | | 3.75 | Figure 5B | -5 | |
| | | 14 | Figure 5C | -50 | |

**Supplementary Table 6. CryoET/EM imaging parameters.** Samples were loaded into one of two cryo-transmission electron microscopes. Two-dimensional images and montages were aligned, and dose-fractionated movies were acquired in counting mode. Tilt series were collected bidirectionally from -21°, through a range from -60° to +60° in 3° increments.

### SUPPLEMENTARY MOVIE LEGENDS

**Supplementary Movie 1. RBS-As are not cylindrical.** An RBS-A partially stuck to a Petri dish fluttered as suction was applied by and released from a micropipette. Note that this video was captured solely to show the morphology of the RBS-As. Debris was inevitably captured as a result of applying sufficient suction for the RBS-A to flutter on its side. No RBS-As were collected for the single-cell genomics experiment during the recording of this video or afterwards using this micropipette.

**Supplementary Movies 2 and 3. CryoET reveals the 3D architecture of RBS-A components.** Yellow, pilus-like appendages; blue, inner membrane; purple, S-layer-like structure; green, outer membrane; red, matrix. Scale bar: 100 nm.

### SUPPLEMENTARY METHODS

Maximum likelihood phylogenies of the Alcaligenaceae family were inferred using the 16S rRNA gene (**Supplementary Fig. 4a**) and ribosomal protein S3 (**Supplementary Fig. 4b**) to confirm the taxonomic affiliation of bin 16, which was recovered from the mini-metagenomics experiment. Gene/protein sequences were acquired for each characterized genus in the Alcaligenaceae, as shown on the NCBI Taxonomy Browser (accessed August 2022), when such sequences were available in the NCBI system (some genera have scant or no genomic representation). We additionally performed BLAST[1] queries of the bin 16 16S rRNA gene and rpS3 protein against the nr/nt and nr databases (accessed August 2022), respectively, and included the top 10 most similar sequences. 16S rRNA gene sequences were aligned using SINA[3] v. 1.2.11, using the SILVA SSU database[4-6] r138.1 as a reference, and columns containing >3% gaps or rows with <50% sequence were removed. rpS3 protein sequences were aligned using Clustal Omega[7,8] v. 1.2.4 and columns containing >5% gaps or rows with <50% sequence were removed. Both phylogenies were inferred using PHYLIP[9] v. 3.1 with 1000 bootstrap replicates, with model selection performed using smart model selection[10] (GTR+R for the 16S rRNA gene and Q.yeast+G+I for the rpS3 protein). Trees were visualized using iTOL[11] v. 6.
